# Joint inference of species histories and gene flow

**DOI:** 10.1101/348391

**Authors:** Nicola F. Müller, Huw A. Ogilvie, Chi Zhang, Michael C. Fontaine, Jorge E. Amaya-Romero, Alexei J. Drummond, Tanja Stadler

**Affiliations:** Fred Hutchinson Cancer Research Center, Vaccine and Infectious Disease Division, 98109 Seattle, USA; ETH Zürich, Department of Biosystems Science and Engineering, 4058 Basel, Switzerland; Swiss Institute of Bioinformatics (SIB), Switzerland; Department of Computer Science, Rice University, Houston TX, USA; Key Laboratory of Vertebrate Evolution and Human Origins, Institute of Vertebrate Paleontology and Paleoanthropology, Chinese Academy of Sciences, Beijing 100044, China; Center for Excellence in Life and Paleoenvironment, Chinese Academy of Sciences, Beijing 100044, China; Laboratoire MIVEGEC (Université de Montpellier, UMR CNRS 5290, IRD 229) et Centre de Recherche en Ecologie et Evolution de la Santé (CREES), Centre IRD de Montpellier, Montpellier, France; Groningen Institute for Evolutionary Life Sciences (GELIFES), University of Groningen, PO Box 11103 CC, Groningen, The Netherlands; Centre for Computational Evolution, University of Auckland, New Zealand

## Abstract

When populations become isolated, members of these populations can diverge genetically over time. This leads to genetic differences between these populations that increase over time if the isolation persists. This process can be counteracted by gene flow, i.e. when genes are exchanged between populations. In order to study the speciation processes when gene flow is present, isolation-with-migration methods have been developed. These methods typically assume that the ranked topology of the species history is already known. However, this is often not the case and the species tree is therefore of interest itself. For the inference of species trees, it is in turn often necessary to assume that there is no gene flow between co-existing species. This assumption, however, can lead to wrongly inferred speciation times and species tree topologies. We here introduce a new method that allows inference of the species tree while explicitly modelling the flow of genes between coexisting species. By using Markov chain Monte Carlo sampling, we co-infer the species tree alongside evolutionary parameters of interest. By using simulations, we show that our newly introduced approach is able to reliably infer the species trees and parameters of the isolation-with-migration model from genetic sequence data. We then use this approach to infer the species history of the mosquitoes from the *Anopheles gambiae* species complex. Accounting for gene flow when inferring the species history suggests a slightly different speciation order and gene flow than previously suggested.

## Introduction

Populations can diverge genetically and become separated over time, due to geography or other factors. Gene flow after populations become genetically isolated can counteract this process (Sousa and Hey, 2013). These events are captured in the genome of sampled individuals of those species. In turn, the genetic sequences of sampled species allow us to draw inferences about their common history (the species phylogeny) by modelling the speciation process. To reconstruct the speciation process from genetic sequence data, the multispecies coalescent model (MSC) can be used (Rannala and Yang, 2003; Liu et al., 2009; Heled and Drummond, 2010). This allows for the reconstruction of the species tree while accounting for discordance between gene trees due to incomplete lineage sorting. Gene flow after populations become genetically isolated can counteract this process of divergence (Sousa and Hey, 2013), and if not accounted for, this can lead to biased inferences about the ancestral history of the species (Leaché et al., 2014). To account for gene flow after speciation, isolation-with-migration (IM) models have been developed (Nielsen and Wakeley, 2001; Hey and Nielsen, 2004; Wilkinson-Herbots, 2008) (see also Sousa and Hey, 2013, for review). Initial Bayesian implementations of the IM model were applicable to only two populations. Further, they could suffer from poor Markov chain Monte Carlo (MCMC) convergence due to an extremely diffuse parameter space (Nielsen and Wakeley, 2001; Hey and Nielsen, 2004).

This difficulty was partly overcome by analytically integrating out some model parameters (population sizes and migration rates) to sample from the posterior distribution of gene trees and speciation times. The gene trees and speciation times are then used to estimate the evolutionary parameters and the effective population sizes and rates of gene flow (Hey and Nielsen, 2007). This approach was later extended to deal with more than two populations (Hey, 2010). All of these methods required the species tree topology and the ordering of speciation times to be known *a priori*.

One of the challenges that restricts joint inference of the species tree and rates of gene flow is that, over the course of an MCMC, different species can co-exist. This means gene flow can happen between different co-existing species at different stages of the MCMC. When operating on the species tree such that the co-existing species change, some of the possible routes of gene flow disappear and some newly appear. At the same time, this means that the migration history of a gene tree, i.e. an explicit sequence of migration events, is possibly no longer valid after a new species tree is proposed. Operating on the species tree is therefore particularly challenging if the migration history of each gene tree has to be explicitly considered as well. In Hey et al. (2018), this challenge was overcome by using a clever mapping of migration events between extant species to ancestral species using what is called a hidden genealogy. This means that for any ranked species history, migration histories are always defined.

Having to infer migration histories can lead to computational issues in the related structured coalescent model (De Maio et al., 2015; Müller et al., 2017). Alternatively, inferring migration histories could be avoided altogether.

Here, we introduce a novel isolation-with-migration model (AIM) that allows joint inference of the species tree with rates of gene flow and effective population sizes that avoids the sampling of migration histories. We do so by extending the marginal approximation of the structured coalescent (Müller et al., 2017) to the isolation-with-migration model. Modelling the movement of lineages between speciation events as a structured coalescent process allows us to evaluate the probability of a gene tree given a species tree, a set of rates of gene flow and effective population sizes. The probability of a gene tree given any species tree and set effective population sizes and rates of gene flow can always be calculated using this framework. Using MCMC sampling, we can then operate on the species tree topology, divergence times, gene trees, rates of gene flow and effective population sizes for extinct and extant species. We implemented this approach as an update to StarBEAST2 (Ogilvie et al., 2017), which is available as a package for the phylogenetic software platform BEAST2 (Bouckaert et al., 2014, 2019).

By using simulations, we show that the AIM model is able to infer rates of gene flow (which are equivalent to migration rates in the structured coalescent model), effective population sizes and species trees reliably from molecular sequences directly. In contrast to AIM, the MSC can strongly support wrong species tree topologies and systematically underestimate speciation times.

We then apply AIM to jointly infer the species trees and the rate of gene flow between individual *anopheles* species from the *Anopheles gambiae* species complex (AGC) (Fontaine et al., 2015). The AGC consists of at least eight distinct species that are morphologically indistinguishable (Davidson, 1962; White et al., 2011). Three of these species are amongst the worlds most important malaria vectors (*An. gambiae, An. coluzzii, and An. arabiensis*). Interestingly, this species complex has become a flagship example of reticulated evolution (Mallet et al., 2016; Clark and Messer, 2015). Deciphering the species tree in the AGC remained a challenge for decades due to the confounding processes of incomplete lineage sorting and gene flow that blurred the species tree. The X chromosome and the autosome of these *anopheles* species have been described to code for vastly different species tree topologies (Fontaine et al., 2015) and only 2% of the genome, mostly located on the X chromosome, has been suggested to reflect the true species order (Fontaine et al., 2015; Thawornwattana et al., 2018). This dataset has been previously analysed using different methods. Fontaine et al. (2015) inferred the speciation history directly from the gene trees themselves, while Thawornwattana et al. (2018) inferred the speciation history by using a multi-species coalescent approach implemented in BPP (Yang, 2015a).

## Results

### Inference of the species tree from genetic sequences

We first test if AIM is able to infer the true species tree, rates of gene flow and effective population sizes of extant and extinct species. To do so, we first simulated 1000 species trees with 4 taxa under the Yule model (Yule, 1925) with a speciation rate randomly sampled from a lognormal distribution with mean=100 and *σ*=0.1. The narrow distribution around the speciation rate is chosen such there are no issues arising from migration rates or effective population sizes being too high or low relative to the species tree.

For each of those 1000 randomly sampled species trees, we next sampled at random the effective population sizes of each species from a lognormal distribution with mean=0.0025 and standard deviation=0.25. Between each co-existing species, we randomly sampled if there is on-going gene flow from a binomial distribution with 5% probability on there being gene flow. This put an approximately 40% probability of there not being any gene flow in the simulated datasets, meaning that these simulations include datasets with and without gene flow. If there was gene flow, we sampled the forward in time migration rate from an exponential distribution with mean=100. Additionally, we say that the rate of each lineage having originated from a different species is at most the inverse time of co-existence between two species. Without that constraint, we would allow for scenarios where speciation events are entirely unobserved.

For each of the 1000 simulated species trees, effective population sizes and migration rates, we simulated 50 gene trees using MCcoal (Yang, 2015b). Each of the four species had 2 sampled individuals. For each of the gene trees, we next simulated genetic sequences using the HKY model with a transition/transversion ratio of 3 and assuming a random relative evolutionary rate scaler drawn from an exponential distribution with mean=1 using SeqGen (Rambaut and Grassly, 1997).

We next used AIM to jointly infer the species tree, rates of gene flow, and effective population sizes of all extant and ancestral species and evolutionary rates from the simulated sequences. For each rate of gene flow between two co-existing species, we estimate the support for this rate to be non zero using the BSSVS approach (Lemey et al., 2009a). As shown in figure 1, effective population sizes, migration rates and speciation times are inferred reliably. As expected, the 95% highest posterior density (HPD) intervals contain the true values under which the simulation was performed in around 95% of all cases.

**Figure 1:**
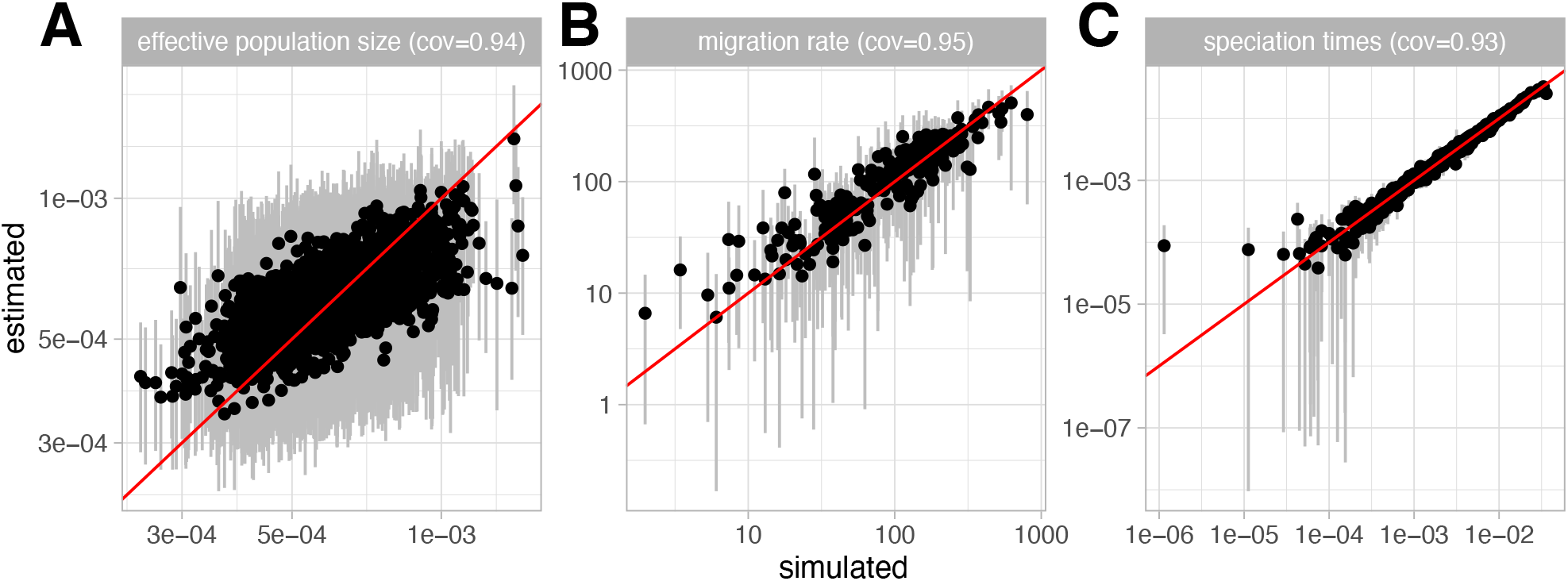
Inference of effective population sizes, migration rates and speciation times. **A** Here we compare estimated effective population sizes on the y-axis to the true simulated effective population sizes on the x-axis. The grey bar represent the 95% highest posterior density interval. **B** Comparison between estimated and simulated migration rates conditional on there being gene flow. The estimated support for gene flow is shown separately in figure 2. **C** Estimated versus simulated speciation times.

**Figure 2:**
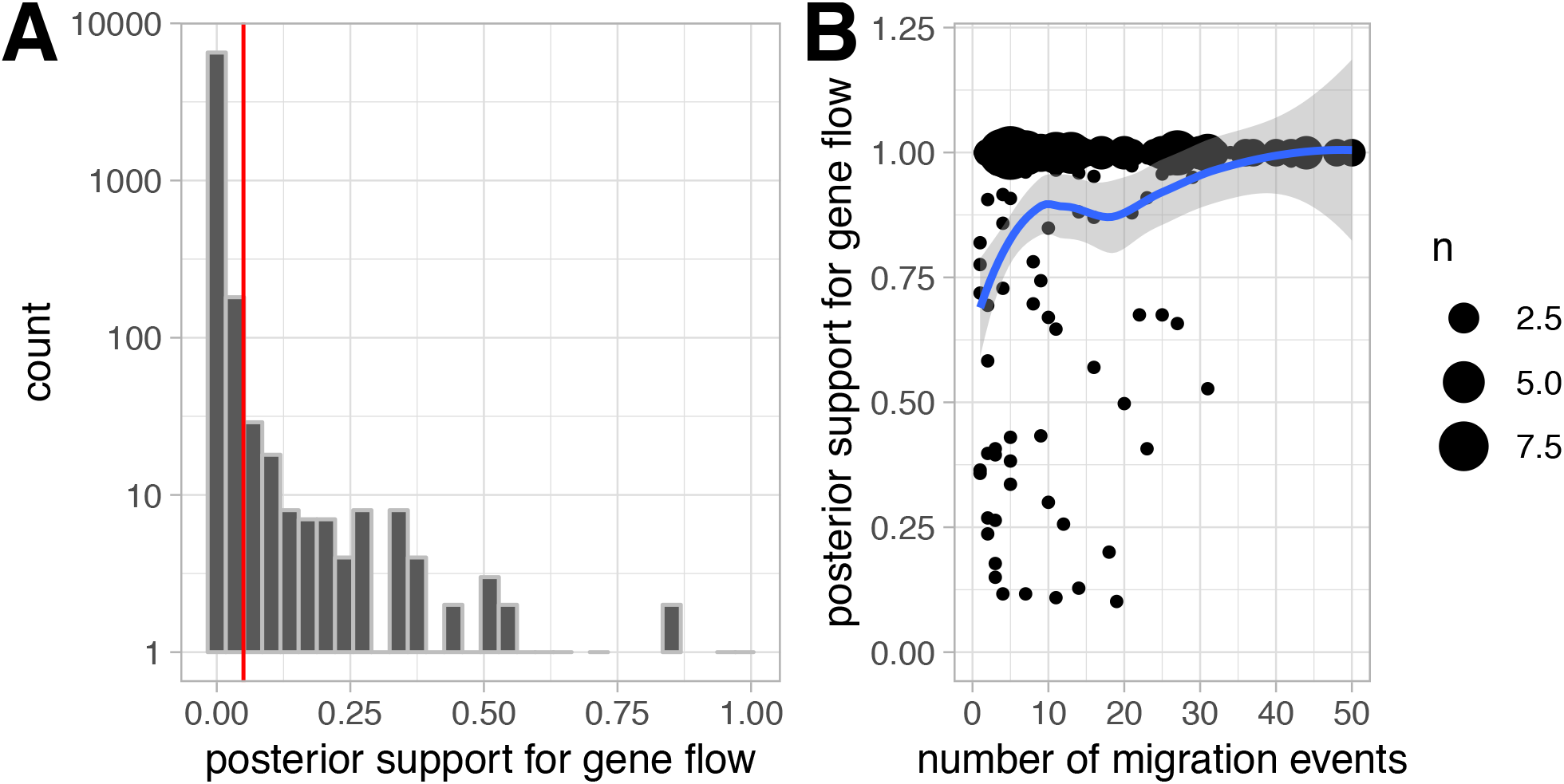
Posterior support for gene flow. **A** Distribution of the support of gene flow between species for which there was no gene flow in the simulations. The x-axis shows the support for gene flow and the y-axis shows the amount of times that posterior support was observed in log scale. The red line shows the prior support for gene flow. **B** Posterior support for gene flow on the y-axis versus the number of migration events between the two species on the x-axis. The curve is a mean estimate for the posterior support of gene flow for different numbers of migration events. The mean estimates are calculated using a loess regression between the number of migration events and the posterior support for gene flow.

We next study the ability of AIM to detect gene flow. To do so, we first computed the distribution of posterior support for gene flow between species for which no gene flow was present in the simulations. As shown in figure 2a, AIM is able to reject gene flow when there is none present. Second, when there are migration events between species, as shown in figure 2b, the model is mostly able to infer that gene flow occurred. This is the case except for a few simulations, where the support for gene flow is lower, but still greater than the prior for gene flow.

Lastly, we compare the inference of species tree topologies and speciation times between accounting for gene flow (AIM) and not accounting for it (MSC). To do so, we analysed the same simulated datasets using StarBeast2 where we jointly infer the species history, effective population sizes of each species as well as all evolutionary parameters. StarBeast2 implements the multispecies coalescent model in BEAST2. For most simulated datasets, both methods infer the species tree topology well, with AIM inferring higher posterior support for the true topology (see figure S1A). The estimates of speciation times are largely consistent between the two methods (see figure S1B).

Between most species, however, there was no gene flow in the simulations. To see when there are differences between the two approaches, we next look at the support for the true species tree topology depending on the overall number of migration events (see figure 3A). With more and more migration events, the support for the true species tree topology decreases using the multi-species coalescent. Using AIM, the support for the true species tree topology is largely independent of the total number of migration events (see figure 3A). Figure 3B shows the estimated minus the true speciation times using the two approaches. These results are shown depending on how many migration events happened between the two clades below each speciation event. The more migration events there were between these two clades, the stronger the underestimation of the speciation time becomes. In other words, if there are migration events between two clades, the multi-species coalescent infers speciation events to have occurred closer to the present. The speciation time estimates using AIM are largely unaffected by this. This observation is consistent with biases in inference of speciation times observed previously (Leaché et al., 2014).

**Figure 3:**
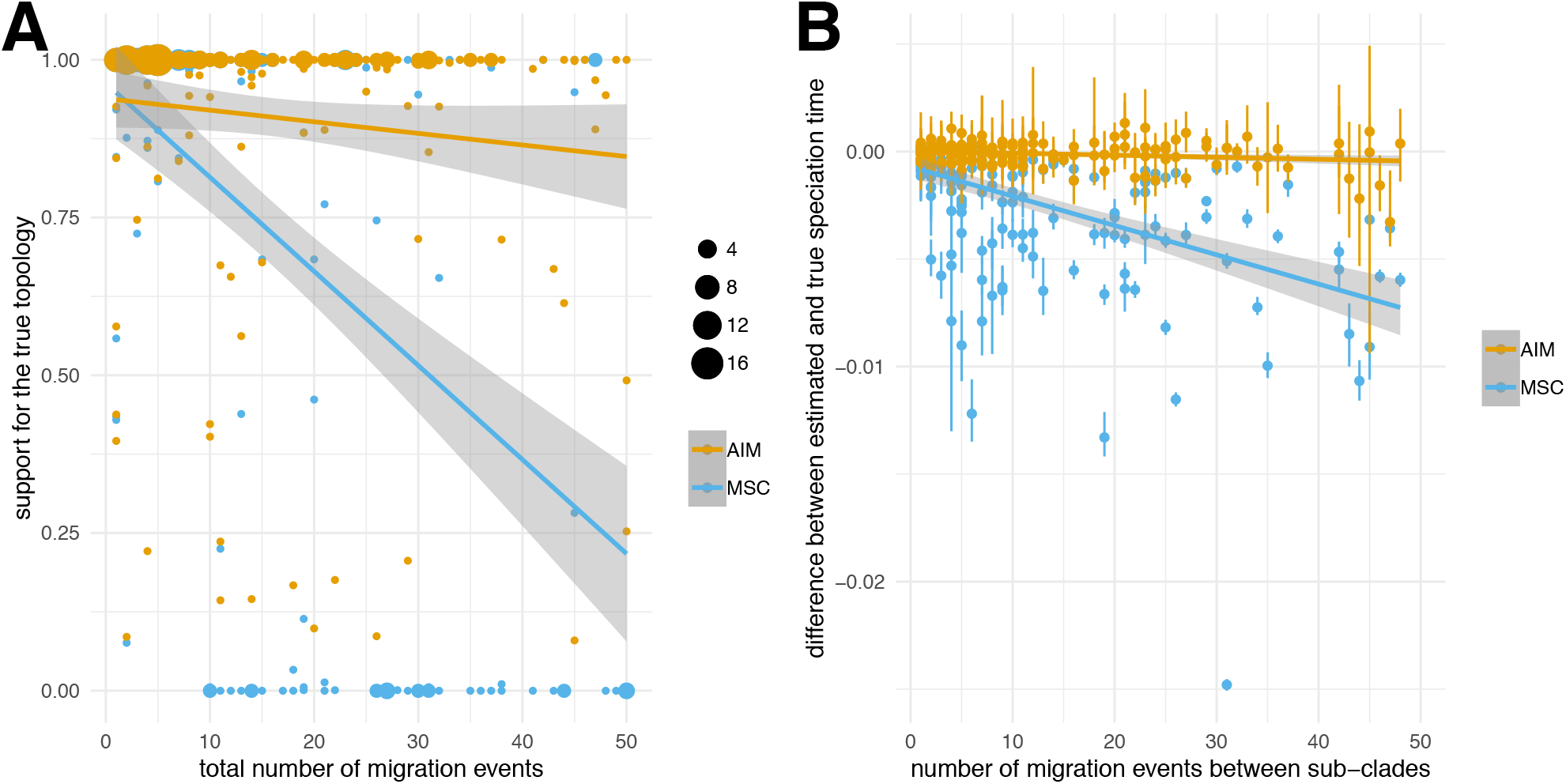
Comparison of species tree and speciation time inference using AIM and MSC. **A** Posterior support for the true species tree topology inferred using the approximate isolation with migration (AIM) model and the multispecies coalescent (MSC). The curve denote the mean support for the true species tree topology calculated using a linear regression between the total number of migration events and the support for the true topology. **B** Comparison between the 95% highest posterior density intervals of speciation times between AIM and MSC. As shown in figure 1C, AIM infers the true speciation times well. The multispecies coalescent is biased towards an underestimation of the speciation time.

### Resolving the evolutionary history of *Anopheles gambiae* complex

Next, we used (AIM) to study the species history of the *An. gambiae* species complex (AGC). Previous studies Fontaine et al. (2015); Thawornwattana et al. (2018) showed that different regions of the genomes of the AGC code for different topologies, especially with respect to the branching of *An. arabiensis*. Most of the X-chromosome (i.e. the Xag inversion) was shown to be indicative of the species branching order, where *An. arabiensis* cluster with *An. quadriannulatus* (Fontaine et al., 2015). In contrast, the autosomes were shown to be strongly impacted by introgressions between *An. arabiensis* and *An. gambiae* or *An. coluzzii* (Fontaine et al., 2015). Here, we assessed the ability of AIM to reconstruct the species history of the AGC in two ways, namely analyzing only the x-chromosome and analyzing both the x-chromosome and chromosome 3. We first split the chromosome into loci of approximately 1000 base pairs.

After removing all loci which had variable sites on more than 50% of all positions, we randomly sampled 200 loci along the X chromosome. In order to control for sensitivity due to this random sub-sampling, we repeated this step 3 times. We then jointly inferred the species history and the support for gene flow between individual species from these regions. We assumed the different regions to evolve according to an HKY+Γ_4_ model with a transition/transversion rate that we estimated for each region individually, while fixing the base frequencies to the observed frequencies. We further allowed each region to have a different relative evolutionary rate. Since we only have loci from one individual per species, we further assumed that the effective population size of all extant species was the same. The reason is that with having only one sampled individual per species, coalescent events in extant species can only occur when there is gene flow to extant species. This means that there are only very few or no coalescent events in a species to inform effective population sizes in extant species.

Figure 4 shows the inferred evolutionary history of 8 anopheles species, including the two outgroup species (*An. christyi* and *An. epiroticus*), averaged over the 3 random subsets. Using the AIM model, we inferred that *An. merus* had its most recent specation event with the common ancestor of *An. coluzzii* and *An. gambiae*. Fontaine et al. (2015) showed that the bottom of the species tree was poorly resolved using classic phylogenetic approaches due primarily to incomplete lineage sorting. In contrast, Thawornwattana et al. (2018) inferred that *An. merus* was the first species to split from the rest of the *An. gambiae* species complex using the MSC and 100 loci. The branching order for the rest of the tree was consistent with both Fontaine et al. (2015) and Thawornwattana et al. (2018).

**Figure 4:**
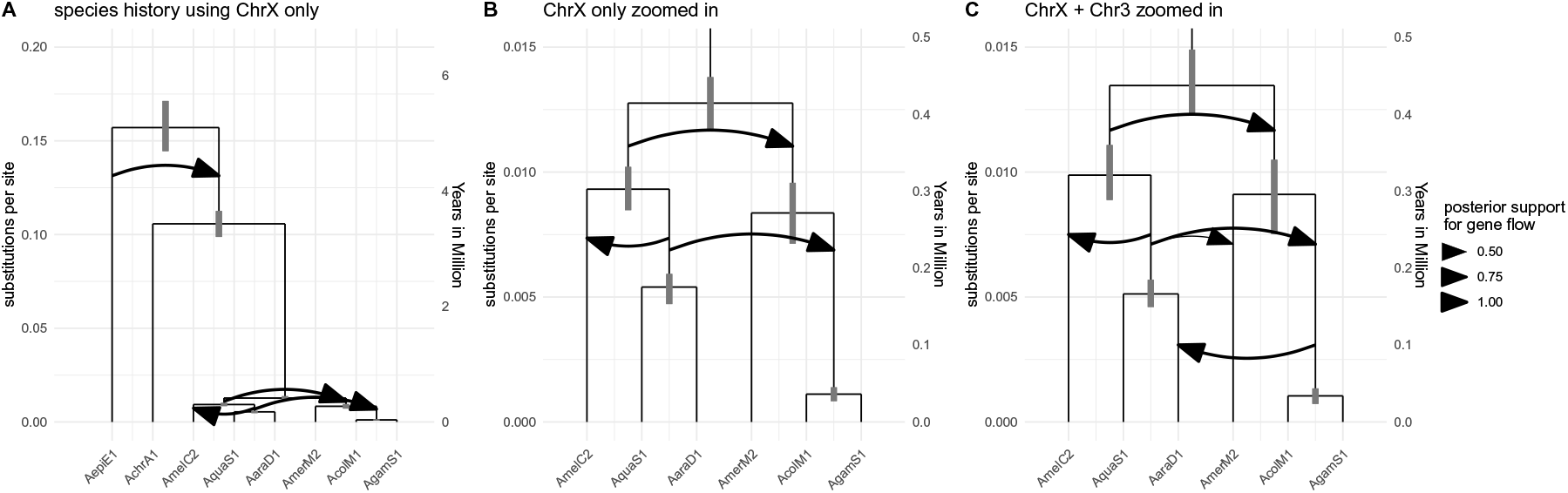
Inferred species evolutionary history of the *An. gambiae* complex. The inferred species history of anopheles is shown in units of substitutions per site averaged over all 3 random subsets of loci from either only the X chromosome (ChrX) or from the X chromosome and chromosome 3 (ChrX + Chr3). The node heights are the median inferred speciation times. The grey bars show the 95% highest posterior density intervals for speciation times. The heights are given in substitutions per site. The cutoff for an arrow to be plotted is support for gene flow with a posterior support of at least 0.5. **A** Inferred species history for all 8 anopheles species including the two outgroups *An. christyi* and *An. epiroticus*, with support for ancestral gene flow between them. This analysis was done by averaging the results over 3 random subsets of 200 loci from the X chromosome. **B** Inferred species history of the same anopheles species but zoomed into the *An. gambiae* species complex. **C** Inferred species history of the anopheles species averaged over 3 random subsets of 200 loci from the X chromsome and chromosome 3. Each of the 200 loci had a 75% chance to be from the X-chromosome and a 12.5% chance to be from the left or right arm of chromosome 3. The inference was done using all 8 species, but the results are zoomed into the *An. gambiae* species complex.

We inferred the common ancestor of all species, expect the outgroups to be about 0.4 Million years ago assuming the same evolutionary rate of 3.08 10^−8^ per year as in Keightley et al. (2014) and Thawornwattana et al. (2018) for non-coding loci. We estimate the isolation events of populations to have happened earlier than the estimates of speciation events in Thawornwattana et al. (2018). Since we, however, sub-sample loci depending on how much variation they have, the speciation time estimates might not be directly comparable. We, for example, also infer a slighly more distant divergence time with *An. christyi* of approximately 0.1 compared to 0.08 substitutions per site in Thawornwattana et al. (2018). If we assume that this difference is due to different sub-sampling of loci, our inferred common ancestor time of the *An. gambiae* species complex is consistent with Thawornwattana et al. (2018).

Jointly with the speciation history, we inferred the presence of gene flow between any co-existing species. Gene flow is indicated by arrows between two co-existing species. Arrows are plotted for gene flow between species with a posterior support of at least 0.5. We find support for gene flow between *An. epiroticus* and the common ancestor of all other species in all random subsets (see figures 4A & S4).

Over all datasets and different number of samples, we infer gene flow from the ancestral species of *An. arabiensis* and *An. quadriannulatus* to the ancestral species of *An. coluzzii* and *An. gambiae*(see figures 4B & S5).

The rates of gene flow were estimated using only loci from the X chromosome, where there is little information about gene flow (Fontaine et al., 2015; Thawornwattana et al., 2018). In fact, most of the information about gene flow has been reported to lie on the autosomes (Fontaine et al., 2015). To test how the inference changes when including loci from the autosomes, we next compiled 3 random datasets of 200 loci with each loci having a 75% chance of being from the X chromosome and a 0.25 and loci from chromsome 3. The higher probability of including loci from the X chromosome is chosen such that there is still enough information about the species tree in the dataset. We then jointly inferred the species tree and gene flow using the same priors and evolutionary models as before for each dataset.

We find support for gene flow between the same co-existing species when including loci from chromosome 3 (see figures 4C & S5).

For all three random subsets, we now find support for gene flow from the ancestral species of *An. coluzzii* and *An. gambiae* to *An. arabiensis*, which is consistent with Fontaine et al. (2015) (see figure S5). This inferred directionality of gene flow is consistent with Fontaine et al. (2015) and Thawornwattana et al. (2018). In 2 of the 3 random subsets, we also find support for gene flow between between *An. merus* and the ancestral species of *An. quadriannulatus* and *An. arabiensis*, but not between *An. merus* and *An. quadriannulatus* as in Fontaine et al. (2015).

## Discussion

The AIM model and its implementation introduced here is able to jointly estimate the species tree topology and times, effective population sizes, and rates of gene flow between species, from multi locus molecular sequence data. These parameters are relevant to many biological systems. Our approach is implemented in a new version of an open source package (StarBEAST2) which is an add-on to the phylogenetics software platform (BEAST2). This means that users of AIM can take advantage of the flexibility of BEAST2, including the large number of available molecular clock models and substitution models.

Using simulations, we demonstrate the validity of our approach as well as the problems that can occur when gene flow is not accounted for. The species tree topologies and node heights are inferred accurately in all scenarios we simulated. Not accounting for gene flow can lead to underestimated speciation times as well as incorrect species tree topologies. This is consistent with previous observations (Leaché et al., 2014). The estimates of rates of gene flow are unbiased but inferring support for gene flow can be complex when only a few gene flow events in the datasets are captured in the gene trees.

When analysing the species history from random loci of the X chromosome of eight *Anopheles* species, we inferred a different speciation order compared to previous results using the multi-species coalescent (Thawornwattana et al., 2018). In particular, we estimate *An. merus* to attach to the common ancestor of *An. coluzzii* and *An. gambia*, whereas Thawornwattana et al. (2018) inferred *An. merus* to be an outgroup to the other species of the ACG complex. This differences can be explained by what the different models consider a speciation event. In the multi-species coalescent, a speciation event is more or less considered the last time genes were exchanged between populations, whereas the isolation-with-migration model considers the initial isolation of populations to be a speciation event.

Jointly with the species history, we inferred gene flow between co-existing species. When only using loci from the X chromosome, we did not find support for gene flow from *An. arabiensis* to the ancestral species of *An. coluzzii* and *An. gambia* as found previously (Fontaine et al., 2015; Thawornwattana et al., 2018). This is expected since genes carrying information about gene flow between those two species are mostly located on the automsomes. Instead, however, we found support for gene flow between the ancestral species of *An. arabiensis* and *An. quadriannulatus* and *An. coluzzii* and *An. gambia*.

When including genes from chromosome 3, we find support for gene flow from *An. arabiensis* to the ancestral species of *An. coluzzii* and *An. gambia* Additionally, we find support for gene flow from the ancestral species of *An. arabiensis* and *An. quadriannulatus* to *An. merus*, but not from *An. quadriannulatus* directly. Random selection of loci could however miss some of the information about gene flow. It remains to be seen if sub-sampling strategies that perform a weighted selection of loci to better reflect the information content across the full chromosome would allow us to better infer gene flow. Additionally, using more loci could allow us to better capture more rare gene flow events. This may, however, require better Markov Chain Monte Carlo operators that are able to more efficiently explore the posterior probability space. In particular, we currently do not utilize Markov Chain Monte Carlo operators that jointly propose changes to gene trees and rates of gene flow between co-existing species. Adding such might substantially increase the amount of loci that can be used for inferences. The implementation of AIM as part of StarBeast2 further allows to include additional sources of data such as fossil data to infer the species tree (Ogilvie et al., 2018) using the Fossilize-Birth-Death model framework (Stadler, 2010). Additionally, since the underlying structured coalescent theory has been developed explicitly for serially sampled data (Müller et al., 2017, 2018), accounting for ancient DNA will be possible in the future. This will mostly mainly require adapting the implementation to allow loci to be sampled through time, analogue to, for example, pathogen sequence data. Additional Potential extensions to the model could only allow migration to occur for a defined period of time after speciation (Wilkinson-Herbots, 2012). Alternatively, additional information, such as about the overlap of habitats of species, could be used to inform gene flow between species in a generalized linear model framework (Lemey et al., 2014; Mueller et al., 2018).

## Materials and Methods

### Calculation of the probability of a gene tree under the approximate isolation-with-migration model

To calculate the probability of a gene tree under the approximate isolation-with-migration (AIM) model, the following probability has to be calculated:

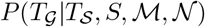

with 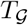 being the gene tree, 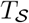 being the species tree, *S* being the species to which each sampled individual belongs 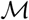 and 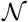 and being the set of migration rates and effective population sizes. To allow for multi-locus data, we assume that each locus evolved independently from the same isolation-with-migration process.

### The location of a gene over time

Isolation-with-migration models are closely related to the structured coalescent process. These models generally require the state or location of every single lineage to be inferred backwards in time. A recently introduced approximation to the structured coalescent avoids this by formally integrating over every possible history (Müller et al., 2017). For each given gene tree 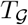, we calculate the probability of a lineage *L_i_* at time *t* being in a particular state *a* (i.e. the gene being in a particular species *a*), 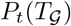. Between speciation on species trees and coalescent events on gene trees, this probability can be described via differential equations as described in Müller et al. (2018) eqn. 1:

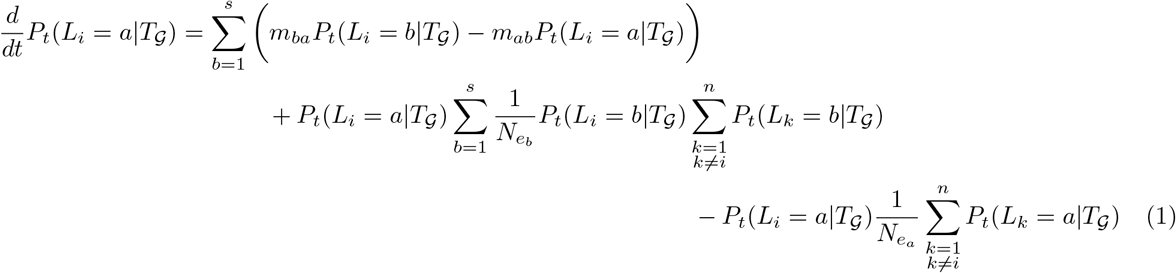

*m_ba_* describes the backwards in time rate at which migration events from species *b* to *a* happen and 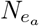 is the effective population size of species *a* and *s* denotes the number of species. At a coalescent event between lineage *i* and *j*, the probability of the parent lineage can be calculated using the following equation (14, page 2979):

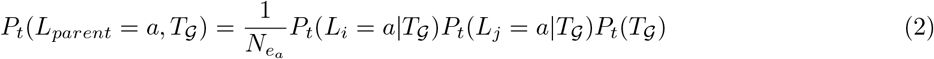

Then we can proceed solving eqn. 1 again backward in time with the initial value 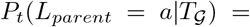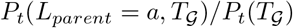

### The location of a gene prior to a speciation event

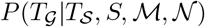 can be calculated similar to the probability of a tree under the structured coalescent. Between speciation events, 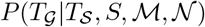 is updated as shown in the previous section. The backwards in time analogue to a speciation event is the combination of two species. If species *a* and species *b* have parent species *c*, the probability of each remaining gene *i* being in species *c* can be calculated as follows:

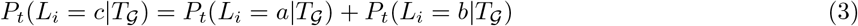

Starting from the present and going back in time, 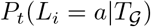 can now be calculated using equations 1 and 2 between speciation events and using equation 3 at speciation events up to the root, 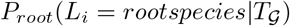.

### Prior assumption of rates of gene flow

We assume the rates of gene flow to be constant over time and we allow them to be either forward or backward in time. Backwards in time rates 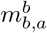 denote the probability of a lineage being in species *b* to have originated from species *a*. The forwards in time rates 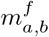 denote the probability of an individual being in species *a* to migrate to species *b*. This rate is not directly accessible in a coalescent framework, but can be approximated as:

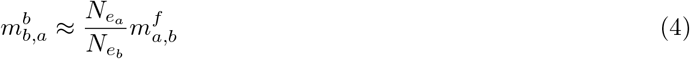

with 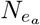 being the effective population size of species *a* and 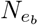 the one of species *b*. This approximation becomes exact if the generation times of both species is the same, that is if 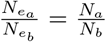. In this manuscript, we always use the forward in time definition of migration rates. Additional to sampling the rate itself, we sample the probability of any migration rate to be 0 by using the BSSVS approach (Lemey et al., 2009b). By sampling any rate being 0 or 1, we can estimate the support for gene flow between two species. Throughout this manuscript, we assume the prior probability for gene flow between two species to be 5%.

Additionally to defining the rates, we implemented the possibility to define maximal rates of migration that depend on the species tree directly. We implemented these in order to allow us to specify a maximal rate of gene flow between two species where we expect that they are essentially not two species. To do so, we implemented two different scenarios:

In the **overlap** scenario, we assume that the maximal backward in time rate of gene flow between two species 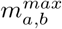 is inversely proportional to the time these species co-exist:

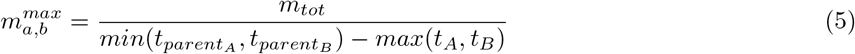

with *t_A_* denoting the node height of *A* and 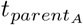 denoting the node height of the parent species of *A*. The variable *m_tot_* denotes an overall rate scaler that can be specified. This allows us to put maximal values on the rate of gene flow that, while not exactly the same, are closely related to the percentage of lineages between these two species to migrate.

In the **distance** scenario, we assume that any the maximal backward in time rate of gene flow between two species 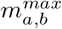 is inversely proportional to the distance between the common ancestor :

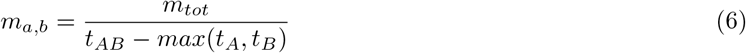

with *t_A_* denoting the node height of *A* and 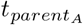 denoting the node height of the parent species of *A*. This is to account for the maximal rate of gene flow being likely smaller between more distant species. Throughout the manuscript, we used the overlap description of rates of gene flow.

The effective population sizes of extant and ancestral species are assumed to be log-normally distributed with *σ* = 0.25. The mean of the log-normal distribution is estimated. Exact specifications for all parameters as well as MCMC operators are provided in the BEAST2 xml input files here https://github.com/nicfel/Isolation-With-Migration.

### Exploring different ranked species tree topologies

In order to operate on the species tree, we use the standard tree operators implemented in BEAST2. Gene flow can only occur between co-existing species. The species which coexist changes when the rank (i.e ordering) of speciation times or the topology of the species tree changes. When this occurs, we keep the rates of gene flow between species that were co-existing before and after the operation the same. Rates of gene flow that disappear during a move are randomly assigned to rates of gene flow between co-existing species that newly appear after a move.

### Summarizing posterior distributions of speciation histories

We implemented two ways to summarize a posterior distribution of species trees. In the first, we distinguish between species trees with the different orderings of speciation events. To summarize the posterior distribution of species trees, we first count the number of unique ranked tree topologies. For each unique ranked tree topology, we then compute the distributions of rates of gene flow, effective population sizes and speciation times. Alternatively, we summarize over posterior distributions of species trees, but ignore the ordering of speciation times. In that case, we compute the distributions of rates of effective population sizes and speciation times. For the rates of gene flow, we only consider those between species that are co-existing in all species trees with the same topology, but potentially different ordering of speciation times.

### Anopheles sequence data

We used the whole genome alignment (WGA) from Fontaine et al. (2015) (see also https://doi.org/10.5061/dryad.f4114) for six species in the *An. gambiae* species complex: *An. gambiae* (AgamS1), *An. coluzzii* (AcolM1), *An. arabiensis* (AaraD1), *An. melas* (AmelC2), *An. merus* (AmerM2), and *An. quadriannulatus* (AquaS1), as well as two Pyretophorus outgroup species (*An. christyi* (AchrA1) and *An. epiroticus* (AepiE1). The *An. gambiae* PEST reference genome (AgamP4) was also included to anchor the chromosome assembled coordinate system, but was not used for any other purpose, given this reference genome is a micture of both *An. gambiae* and *An. coluzzii*. In Fontaine et al. (2015), two WGA’s were generated. Here we used the WGA generated based on the reference assembly for each species using the MULTIZ feature from the Threaded Blockset Aligner package v.12 (Blanchette et al., 2004).

Based on the *AgamP4* PEST coordinate system, the WGA is partitioned into five chromosome arms: 2L, 2R, 3L, 3R, and X (the unplaced, draft Y, and mitogenome were not considered here). The AIM approach assumes free recombination among loci, but no recombination within a locus. To meet those assumptions, we subdivided the chromosome into loci of 1000 base pairs, a length small enough to minimize the probability that recombination occurred within loci (Thawornwattana et al., 2018). (Thawornwattana et al., 2018) noticed that local realignment was required to fix some misalignment in the original TBA WGA of Fontaine *et al.* (2015). Thus we realigned all loci using MAFFT v.7.394 (Katoh and Standley 2013), using the iterative refinement method (the L-INS-i option), following (Thawornwattana et al., 2018). Only loci from the non-coding portion of the genome were selected for further analyses, following the gross assumption that these loci would be closer to neutrality than the coding regions.

We additionally removed loci that either badly aligned or were too divergent, which can also be a sign of aligning badly. To do so, we only used loci in the analysis where at most 20% of all positions were gaps and at most 40% of positions had nucleotide variations.

We assumed the genetic sequences to evolve according to an HKY+Γ_4_ model with a transition/transversion rate that we estimated for each region individually, while fixing the base frequencies to the observed frequencies. Additionally, we allowed each locus to have its own relative rate of evolution. Finally, we randomly subsampled 200 loci from the X chromosome and then jointly inferred the speciation history, gene flow, effective population sizes and all evolutionary parameters. Additionally, we generated datasets with ≈ 5% from the left arm of chromosome 3, ≈ 5% of the right arm of chromosome 3 and the rest from the X chromosome. We repeated these step 3 times, in order to have 3 random subsets of loci and then ran each analyses using 2 different priors on the migration rates. We additionally repeated all analysis using only 50 loci, as in Thawornwattana et al. (2018), instead of 200 loci.

All the manipulations and processing of the WGA (in MAFF format) were conducted using Maffilter 1.3 (Dutheil et al., 2014).

### Software and Data Availability

Simulation of gene trees given a species tree and migration rates were performed using the software MC-coal in BPP (Yang, 2015b). Simulations of genetic sequences of length 1000 were performed using Seq-Gen 1.3.3 (Rambaut and Grassly, 1997), using a HKY site model with a transition/transversion ratio of 3 and base frequency of 0.3,0.2,0.2 and 0.3. Data analyses were performed using BEAST 2.5 (Bouckaert et al., 2019). The analysis of the *An. gambiae* species complex was performed using parallel tempering in the coupled MCMC package (Mueller and Bouckaert, 2019). The source code of the BEAST2 package AIM can be downloaded here: https://github.com/genomescale/starbeast2. All the scripts used in this study are publicly available at https://github.com/nicfel/Isolation-With-Migration. Analyses were done using Matlab R2015b. Plotting was done in R 3.2.3 using ggplot2 (Wickham, 2009). Trees were analysed by using ape 3.4 (Paradis et al., 2004) and phytools 0.5-10 (Revell, 2012). A tutorial on how to set-up an AIM analysis can be found through the https://taming-the-beast.org/ platform (Barido-Sottani et al., 2017).

## Acknowledgement

NM and TS were funded in part by the Swiss National Science foundation (SNF; grant number CR32I3 166258). CZ is supported by the 100 Young Talents Program of Chinese Academy of Sciences and the Strategic Priority Research Program of Chinese Academy of Sciences (XDB26000000). JEAR was supported by a PhD fellowship from the GELIFES Adaptive Life program of the University of Groningen (The Netherlands).

## Supplementary Material

**Figure S1:**
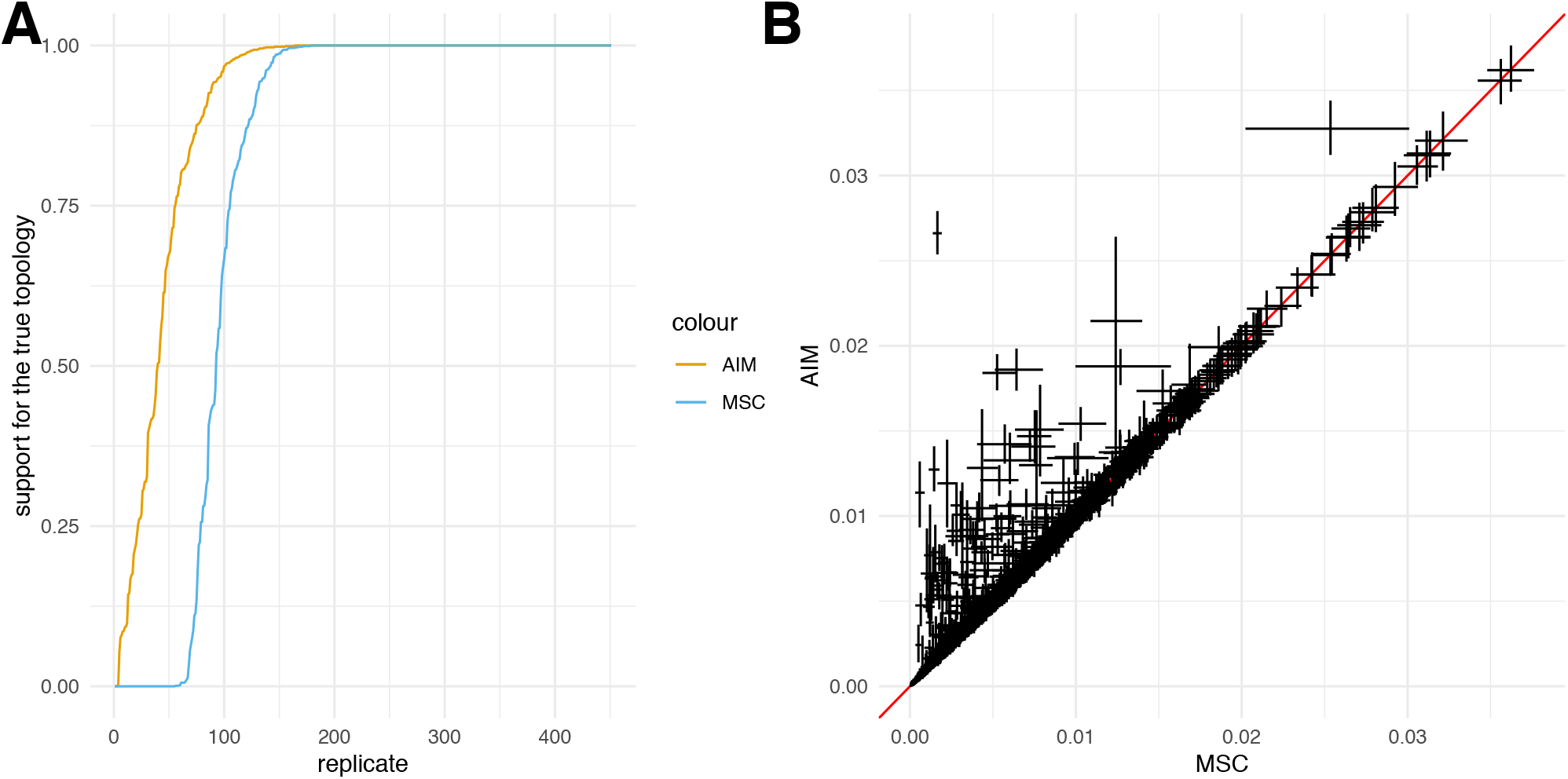
Comparison of species tree and speciation time inference using AIM and MSC. **A** Posterior support for the true species tree topology inferred using the approximate isolation with migration (AIM) model and the multispecies coalescent (MSC) **B** Comparison between the 95% highest posterior density intervals of speciation times between AIM and MSC. As shown in figure 1C, AIM infers the true speciation times well. The multispecies coalescent either correctly infers speciation times or underestimates them.

**Figure S2:**
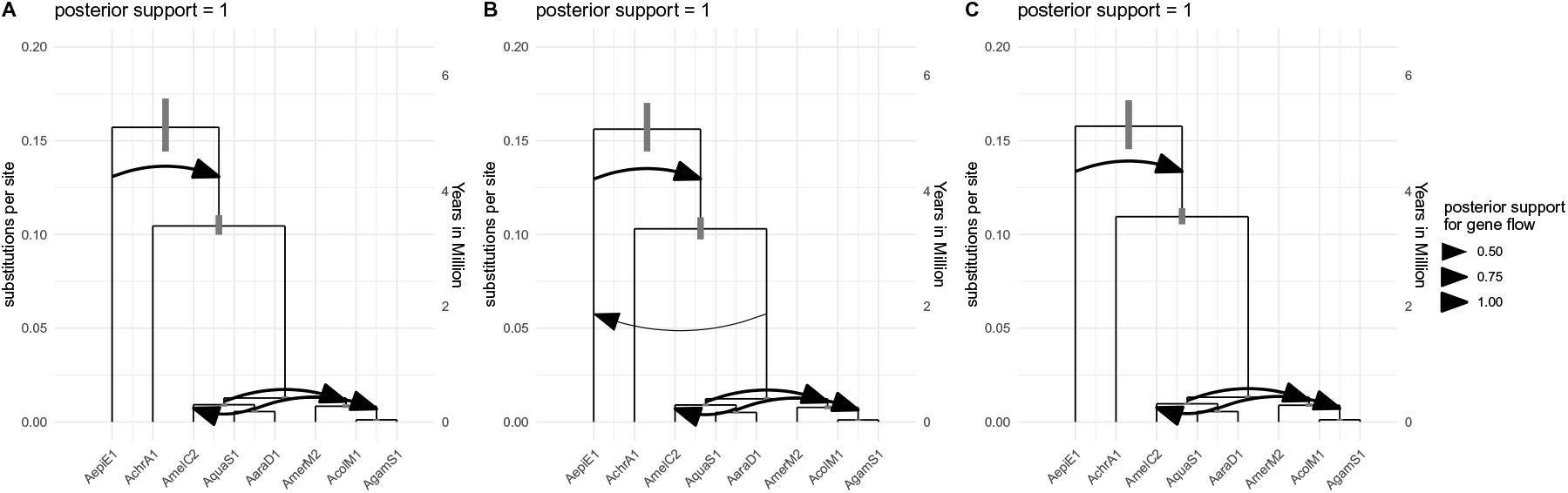
Inferred species histories with gene flow for 3 random subsets of 200 loci from the X-chromosome. Inferred species history for all 8 anopheles species including the two outgroups *An. christyi* and *An. epiroticus*, with support for ancestral gene flow between them. The 3 different trees show the results for 3 different subsets of 200 loci randomly sampled from the X-Chromosome. The node heights are the median inferred speciation times. The grey bars show the 95% highest posterior density intervals for speciation times. The heights are given in substitutions per site. The cutoff for an arrow to be plotted is a posterior support for gene flow of at least 0.5.

**Figure S3:**
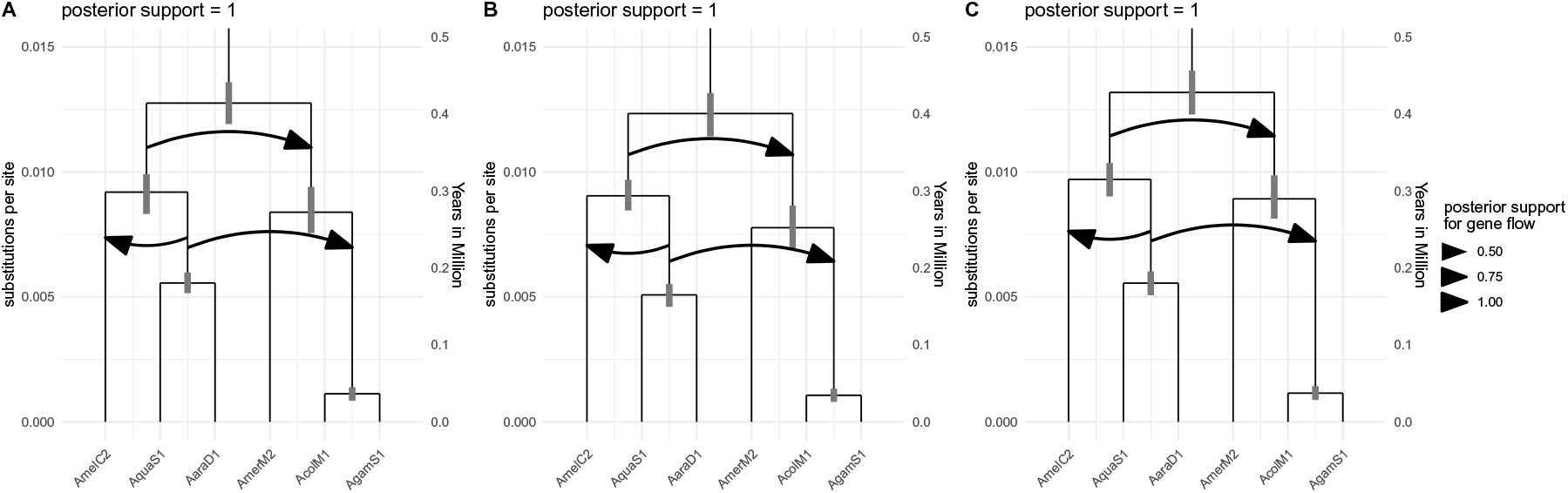
Inferred species histories with gene flow for 3 random subsets of 200 loci from the X-chromosome zooming in into the *An. gambia* species complex. Inferred species history for all 6 anopheles species without the two outgroups *An. christyi* and *An. epiroticus*. The node heights are the median inferred speciation times. The grey bars show the 95% highest posterior density intervals for speciation times. The heights are given in substitutions per site. The cutoff for an arrow to be plotted is a posterior support for gene flow of at least 0.5.

**Figure S4:**
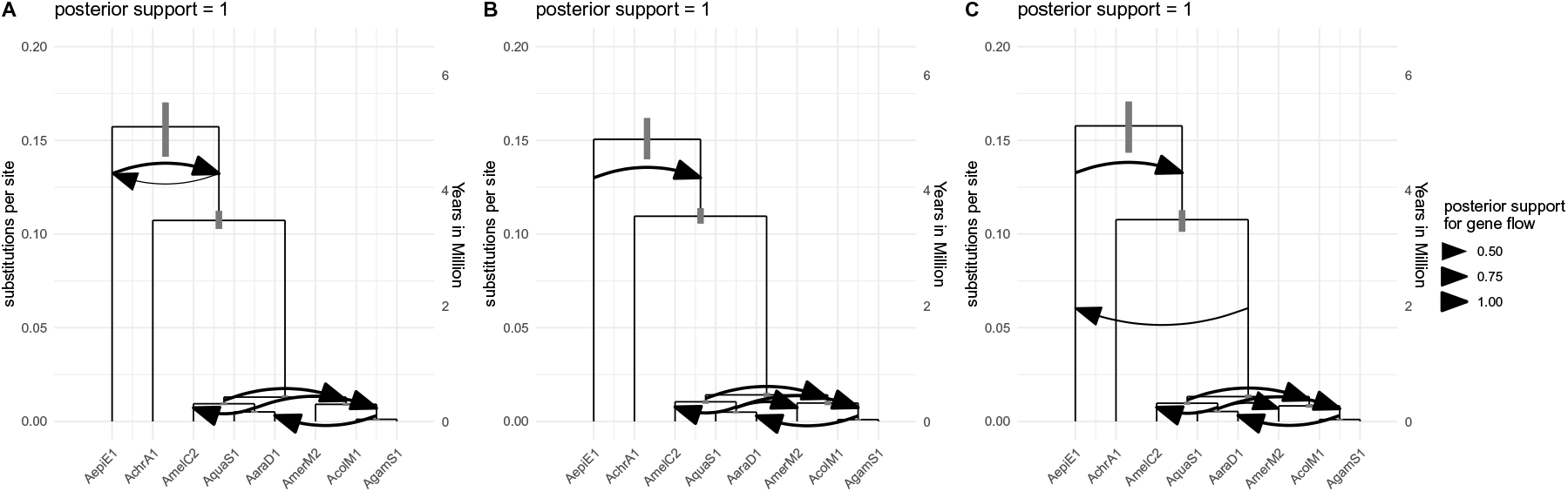
Inferred species histories with gene flow for 3 random subsets of 200 loci from the X chromosome and chromosome 3. Inferred species history for all 8 anopheles species including the two outgroups *An. christyi* and *An. epiroticus*, with support for ancestral gene flow between them. The 3 different trees show the results for 3 different subsets of 200 loci randomly sampled from the X chromosome (with probability 0.9) and chromosome 3 (with probability 0.1). The node heights are the median inferred speciation times. The grey bars show the 95% highest posterior density intervals for speciation times. The heights are given in substitutions per site. The cutoff for an arrow to be plotted is a posterior support for gene flow of at least 0.5.

**Figure S5:**
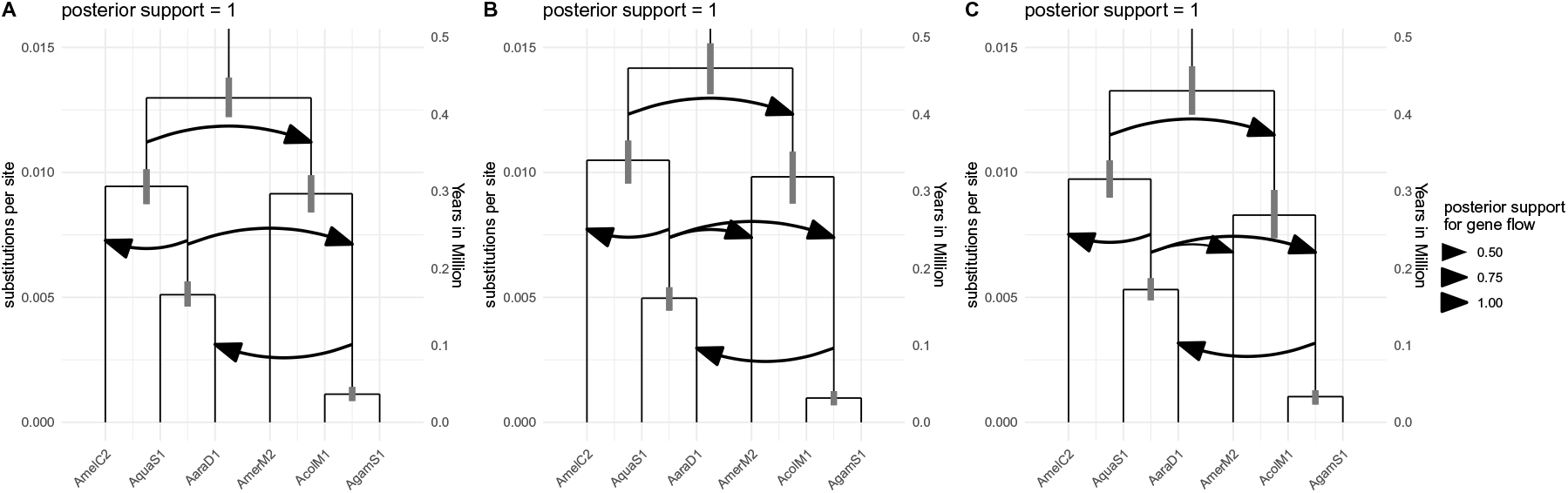
Inferred species histories with gene flow for 3 random subsets of 200 loci from the X-chromosome zooming in into the *An. gambia* species complex. Inferred species history for all 6 anopheles species without the two outgroups *An. christyi* and *An. epiroticus*. The node heights are the median inferred speciation times. The grey bars show the 95% highest posterior density intervals for speciation times. The heights are given in substitutions per site. The cutoff for an arrow to be plotted is a posterior support for gene flow of at least 0.5.

